# SVAPLSseq: A Method to correct for hidden sources of variability in differential gene expression studies based on RNAseq data

**DOI:** 10.1101/062125

**Authors:** Sutirtha Chakraborty

## Abstract

RNAseq technology has revolutionized the face of gene expression profiling by generating read count data measuring the transcript abundances for each queried gene. But on the other side, the underlying technical artefacts generate a wide variety of hidden effects that may potentially distort the primary signals of differential expression between two sample groups. This is in addition to the factors of unwanted biological variability may give rise to a highly complicated pattern of residual expression heterogeneity in the data. Standard normalization techniques fail to correct for these latent variables and leads to a substantial reduction in the power of common statistical tests for differential expression. Here I introduce a novel method SVAPLSseq that aims to capture the traces of hidden variability in the data and incorporate them in a regression framework to re-estimate the primary signals for finding the truly positive genes. Application on both simulated and real-life RNAseq data shows the superior performance of the method compared to other available techniques. The method is provided as an R package ‘SVAPLSseq’ that has been submitted to Bioconductor.

## Introduction

Latent variability is a very important source of residual heterogeneity in gene expression studies. Several latent effects stemming from a wide array of hidden/unknown factors can distort the actual gene expression signals of the tissue samples and introduce spurious sources of extraneous variation. A small number of research works have been conducted over the past few years to address this problem with limited success. Moreover, with the advent of the RNA sequencing technology quantification of gene expression profiles is now possible at a deeper and finer resolution. But the complexity of the RNAseq workflow gives rise to a number of technical artifacts and other potential latent variables that add to the underlying variability in the data (Marioni et al. 2008; Mclntyre et al. 2011). Moreover, the discrete nature of the data hinders the application of standard probability models and necessitates a more generalized approach to analyse it. Methods like SVA (Leek and Storey 2007; Leek 2015), PCA and RUV (Risso et al., 2014) have been developed that have been shown to work effectively for certain specific types of latent variation, while they fail to capture a large number of other unaccountable biases in gene expression data. Recently Leek (2015) developed an extension of the original SVA methodology (SVASEQ) to correct for hidden variability in RNAseq data on gene expression. Although the method conceptually resembles the earlier version, a new approach has been introduced in SVASEQ that uses the information on control probes to extract the signatures of residual heterogeneity. RUV uses a factor analysis approach to identify the sources of latent variability in the gene expression data. Overall, SVA and RUV are robust methodologies that can handle simple latent effects with considerable success, but fails to improve the detection power of differential expression studies in case the hidden variability is highly complicated in nature.

## The SVAPLSseq Methodology

Let *Y_ijk_* denote the read count value measuring the expression level of gene *i* in sample (replicate) *k* from group *j* (*i* = 1(1)*n*; *k* ∈ *S_j_*;*j* = 1,2). Here *S_j_* denotes the collection of samples/replicates corresponding to group *j* (*j* = 1, 2). Let *m* denote the total number of samples in the two groups.

Initially a regression model is fitted on a transformed version of the read count data on gene expression with the sample group (tissue type) as covariate:

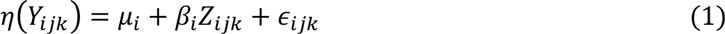

Where *η* is the transforming function, usually chosen as (*x*) = log_e_(*x* + *c*), *c* being a constant, *μ_i_* is the intercept term (baseline effect) for gene *i*; *β_i_* is the regression coefficient for the group indicator *Z_ijk_* (= 1 *if k* ∈ *S*_1_,= 0 *if k* ∉ *S*_1_) corresponding to gene *i* and *ε_ijk_* is the random error term.

Now, residuals from the above fit 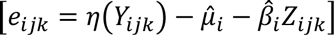 and the transformed read count values [*η(Y_ijk_)*] for all the genes are column-wise organized into two *m* × *n* matrices *E* and *Y* respectively. To that end, *E* and *Y* can be characterized as two *n* dimensional random variables where each dimension corresponds a certain gene. The rationale of the method is to integrate the residual effects in *E* and the original expression signals in *Y* in order to extract the potential signatures of hidden (latent) variability in the data. The multivariate Non-linear Partial Least Squares (NPLS) regression technique is used for this purpose (Boulesteix and Strimmer 2007).The NPLS model for regressing *E* on *Y* is given by:

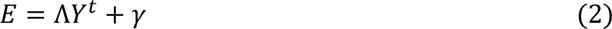

where Λ is the matrix of regression coefficients and γ is the random error component in the model. Now the coefficients in Λ are estimated by partial least squares based on the following two latent factor models for *E* and *Y*

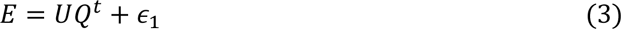

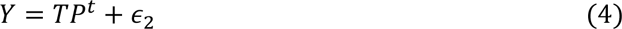

Here *U* = [*u*_1_:*u*_2_:…:*u*_*f*_] is a matrix of *f* unknown, arbitrary latent factors in the column space of *E* and *Q* = [*q*_1_:*q*_2_:…:*q_f_*] is the corresponding matrix of factor loadings.

Similarly *T* = [*t*_1_:*t*_2_:…:*t_f_*] and *P* = [*p*_1_:*p*_2_:…:*p_f_*] are the factor and loading matrices in the column space of *Y*. *ϵ_1_* and *ϵ_2_* are the random error terms in the two models. The algorithm aims to estimate the latent factor pairs (scores) [(*u_i_, t_i_*), *i* = 1(1) *f*] in such a way that their mutual covariance is maximized.

The estimated *Y*-space scores *t*_1_,*t*_2_… ‥ *t_f_* can be visualized as surrogate variables capturing potential signatures of latent variation in the data. These surrogate variables are further tested for significance using a statistical test on their coefficients from the following linear model:

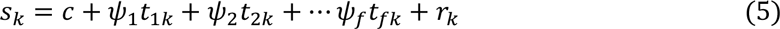

Where *s_k_* is the *k*th component of the first eigenvector of *E^t^E, c* is the intercept, *Ψ_u_* is the regression coefficient for *t_u_* (*u* = 1,2‥ *f*) and *r_k_* is the random error term in the model.

I also introduce another variant of the SVAPLSseq methodology (supervised-SVAPLSseq) which can be used to adjust for the hidden sources of variation in the data when information on control probes is provided. This variant uses the following NPLS regression model for extracting the signatures of latent variability in the data:

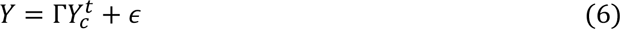

where, *Y_c_* is a matrix of transformed gene level read counts for a predefined set of control genes. Furthermore, two separate regression models are used for *Y* and *Y_c_*, which characterize the latent factors in the column spaces of the two matrices. Similar to unsupervised version of the method, here also the surrogate variables are defined as the PLS scores in the covariate space of the regression model (column space of *Y*_c_). These scores being generated from the vector space of the control gene expression matrix are expected to be free from the primary signals of differential expression and only contain signatures of the unknown variability in the data. Similar to the unsupervised version here also the surrogate variables are tested for statistical significance using an analogous linear model as in (5), with *s_k_* being the *k*th component of the first eigenvector of 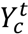*Y_c_*.

The statistically significant surrogate variables can now be incorporated in a regression model and the group effects for all the genes can be re-estimated with higher accuracy. Using these variables in consort with a statistical test for differential gene expression can substantially improve the detection power and specificity of the analysis.

## Simulation Study

The application of the method SVAPLSseq has been illustrated on a simulated RNAseq dataset affected by hidden variables stemming from multiple experimental and sample specific artefacts.

Suppose an RNAseq experiment has been performed to measure the expression levels of 1000 genes over 20 samples distributed equally between two different tissue types. Let *Y_ijk_* denote the number of reads corresponding to gene *i* in sample *k* belonging to the group *j* (*i* = 1(1)1000;*k* ∈ *S_j_*; *j* = 1,2);*S_1_* = {1,2,…10},*S*_2_ = {11,12,… 20}. Here *Y_ijk_* is generated as: *Y_ijk_* = *Z_ijk_* + *h_ijk_* where *Z_ijk_* follows a negative binomial *NB*(*μ_ijk_*, *ϕ_ijk_*) distribution with mean *μ_ijk_* and size (dispersion) parameter *ϕ_ijk_*, *h_ijk_* is the latent effect from the hidden variability in the data.

The gene expression values are assumed to be distorted by multiple sources of hidden variability attributed to several technical artefacts. To that end, let us assume that the 20 samples have been sequenced with two different library preparations, in 4 flowcells (with 5 lanes in each flowcell). Samples indexed (1, 2…10) are sequenced in library 1 and (11, 12…20) are sequenced in library 2. Thus the library effect here is a complex latent variable that is strongly confounded with the primary group effect. Additionally, the samples indexed (1, 2…5) are processed in flowcell 1, (6, 7…10) in flowcell 2, (11, 12…15) in flowcell 3 and (16, 17…20) in flowcell 4. Additionally, it is also assumed that the GC content affects the read counts for each gene. We define 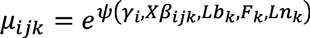, *γ_i_* being the mean parameter for gene *i*, estimated from the zebrafish RNAseq data by using the R package *polyester* (Frazee et al. 2015). *X* is the design matrix for the primary signal of group-specific differential expression and *β_ijk_*~*N*(0,1) *if i* ≤ 400; =0 *if i* > 400. *Lb_k_* is the library effect, *F_k_* is the flowcell effect and *Ln_k_* is the lane effect corresponding to sample *k* and *ΨO* is a continuous function. The library effect *Lb_k_* is generated from an uniform *U*(−0.5,0) distribution if sample *k* ∈ *Library 1* and an uniform *U*(0.7,1) distribution if sample *k* ∈ *Library 2*. The flowcell effect is assumed to be a fixed effect and is generated as:

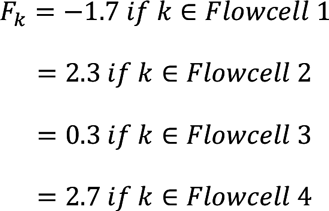

The corresponding lane effect *Ln_k_* for sample *k* in flowcell *u* is simulated from a normal *N(F_U_,* 0.05) distribution with mean *F_u_* and standard deviation 0.05 (*u* = 1,2,3,4). Here, *h_ijk_* is the GC-content effect due to the ith gene on the *kth* sample belonging to group *j* and is generated from a *Binomial* (500, *p_i_*) distribution, *p_i_* being the corresponding GC-content proportion of the gene. A heatmap of the simulated data is presented in Figure 1.

**Figure 1.**
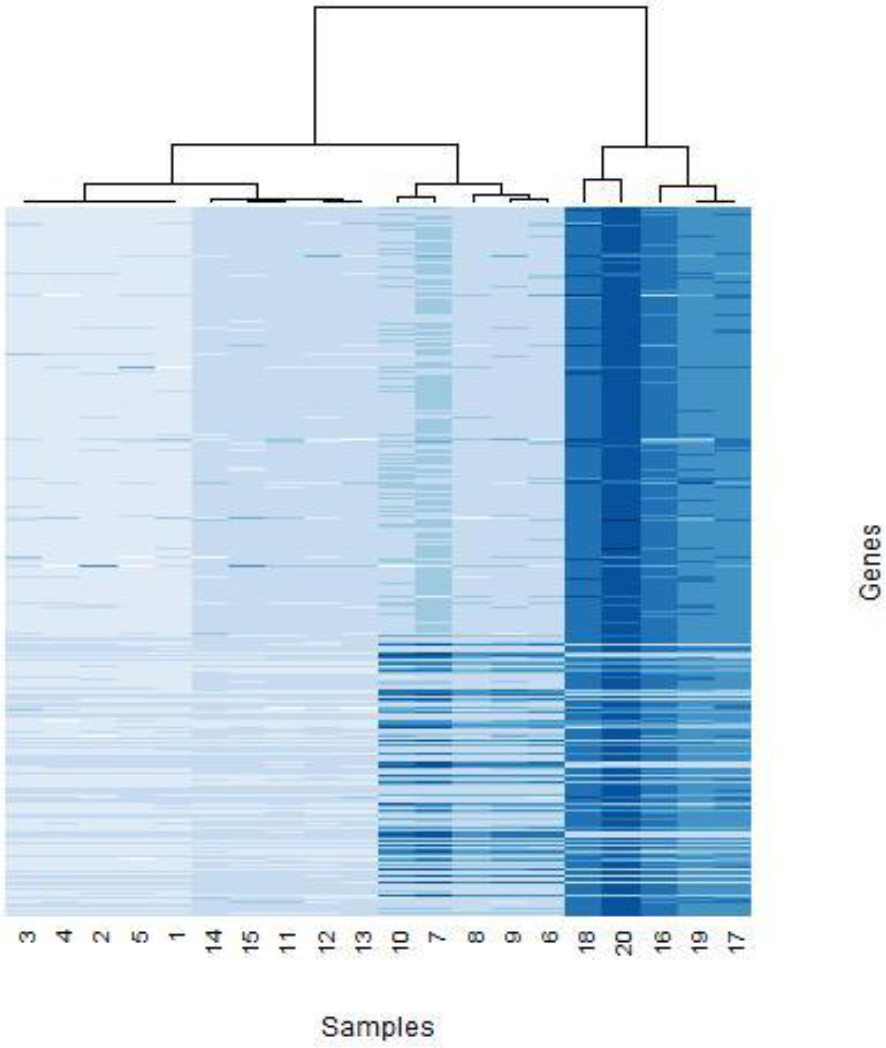
A heatmap of the simulated RNAseq expression data on 1000 genes, showing a strong prominence of the different hidden effects.

The simulation study is based on a performance analysis of the six methods: SVAPLSseq, SVA-IRW, SVA-TWOSTEP, PCA, RUV and standard linear regression on the simulated RNAseq data affected by the complicated pattern of hidden variation. The methods are used to extract the signatures of hidden variability in the data that are in turn used in a linear regression model for estimating the primary signals of differential gene expression. The R package “limma” (Ritchie et al. 2015) is used to fit the hidden effect adjusted models for the different genes and reestimate the primary signals.

The detection power (sensitivity) values from the five methods are presented in Figure 2. Both the variants of SVAPLSseq detect the truly differentially expressed genes with the higher power and accuracy compared to the other methods. In addition, I also computed the correlation coefficients between the simulated latent variables corresponding to the flowcell, lane, library, GC-content effects and their estimated signatures from the different methods. A graphical representation of the corresponding correlation patterns (Figure 3) demonstrates the superiority of SVAPLSseq in accurately capturing the different factors of hidden variation in the data. Higher correlations of the simulated and estimated effects are observed for SVAPLSseq compared to the other methods. Particularly, the correlation profile for SVAPLSseq is strongest for the flowcell and library effects followed by the lane and GC-content effects. This is expected as the first two effects are fixed with respect to the genes while the last two effects vary randomly with each gene.

**Figure 2.**
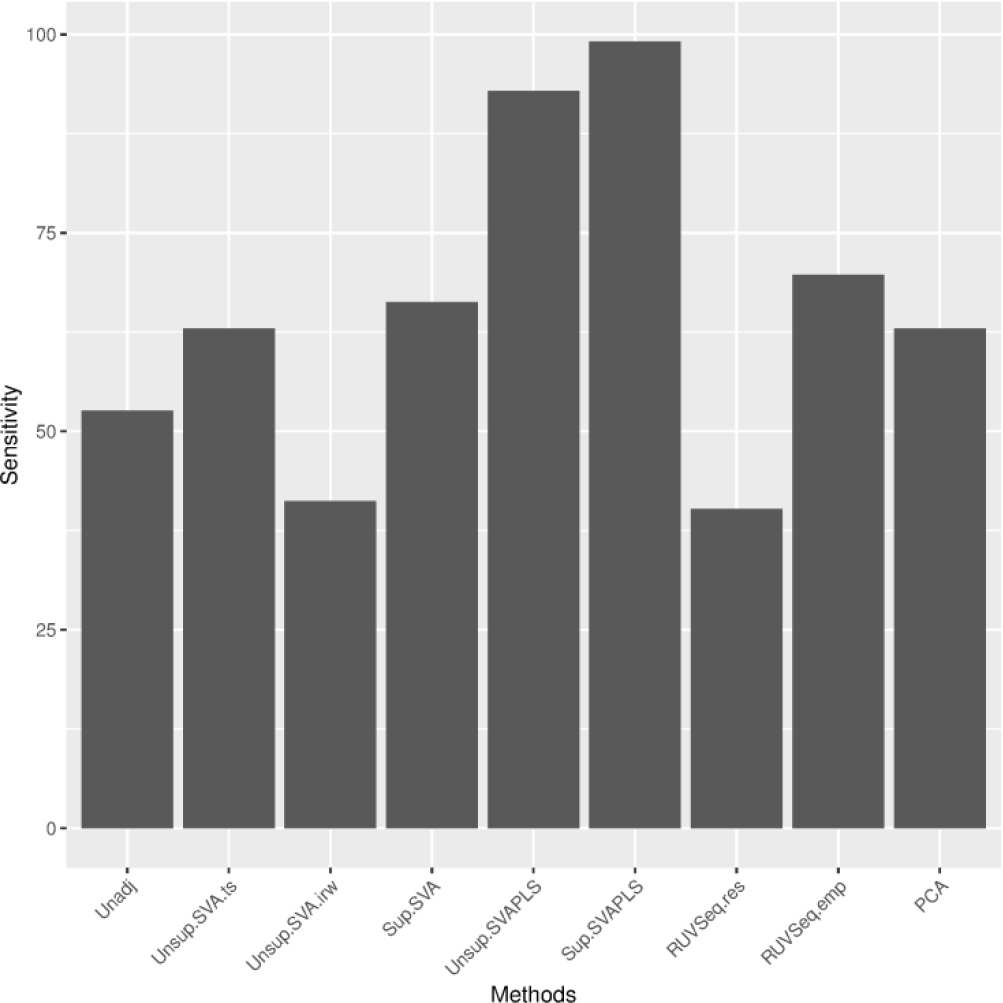
A Barplot showing the sensitivity (detection power) of the different methods on the simulated RNAseq data affected by several hidden variables.

**Figure 3.**
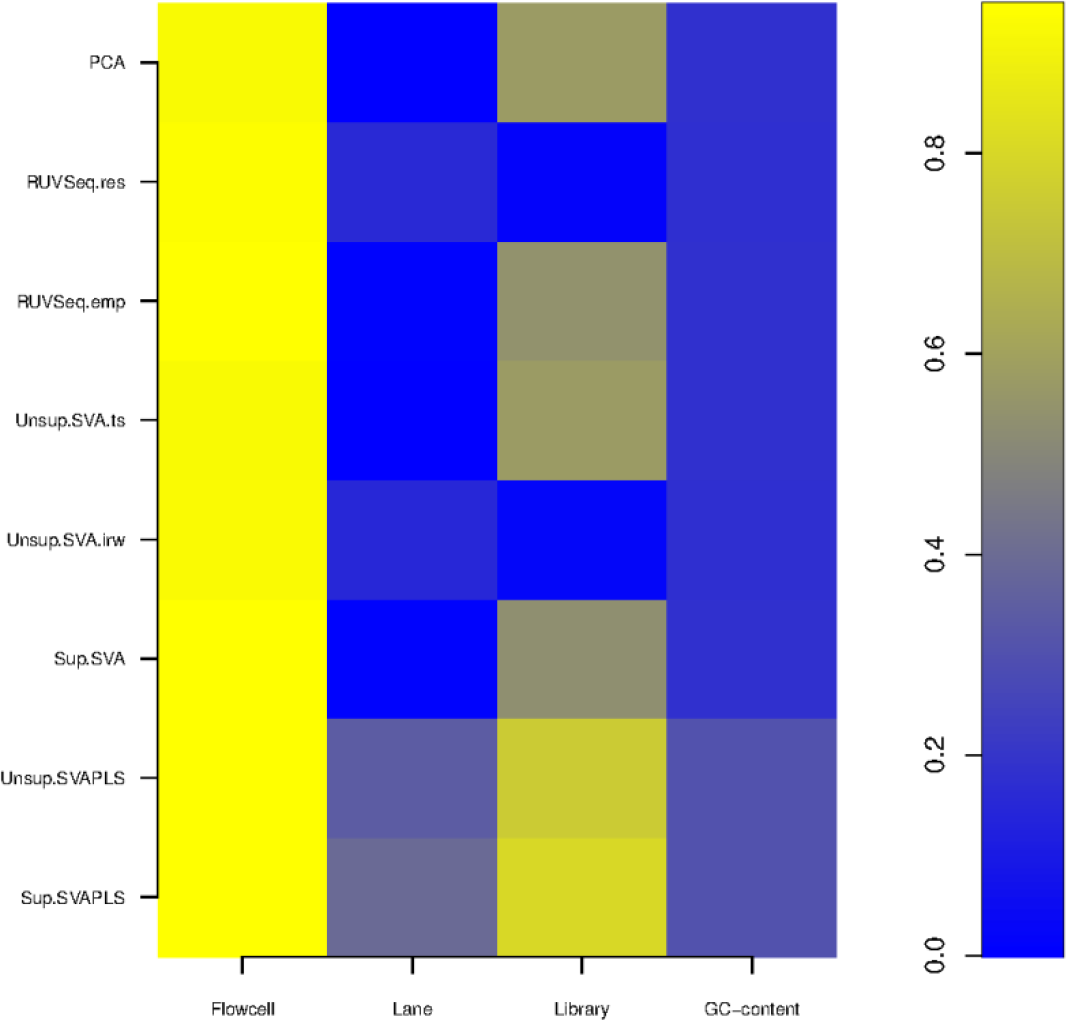
An image plot showing the absolute correlation patterns of the simulated hidden effects and their corresponding signatures estimated from the different methods.

## Application to Real Life Data

In this section a real life application of the methodology has been illustrated on a RNAseq gene expression data from the C57BL/6J (B6) and DBA/2J (D2) strains of 20 inbred mice (Bottomly et al. 2011). The data contains the expression levels of 36356 genes over 20 mice brain samples (11 from B6 and 10 from D2 strain). The data has been trimmed so that every gene has at least one read from a sample in each group. The reduced data has information on 12839 genes. Now, the samples have been sequenced via three different experiments. Distribution of the samples among the three experiments is shown in Table 1. A heatmap of a subset of the data shows a prominent batch effect due to the experiment number (Figure 4). I applied SVAPLSseq on this data along with the five other competing methods in order to adjust for this unwanted variability and detect the differentially expressed genes between the two strains. Overall, SVAPLSseq detected 1713 genes with 1103 genes from PCA, 1796 genes from SVA and 1997 genes from RUVSeq (Figure 5). I also performed a principal component analysis of the residual data obtained after removing the primary signals and the estimated signatures of hidden variability from the *6* methods. SVAPLSseq and SVA exhibited the best performance by accurately adjusting for the inherent batch effect in the data (Figure 6)

**Table 1.** Table showing the distribution of the samples among the three experiments in the Bottomly RNAseq data.

| Experiment Number | Sample Numbers |
| --- | --- |
| 4 | 8, 9, 10, 11, 12, 13, 14 |
| 6 | 1, 3, 5, 7, 16, 18, 20 |
| 7 | 2, 4, 6, 15, 17, 19, 21 |

**Figure 4.**
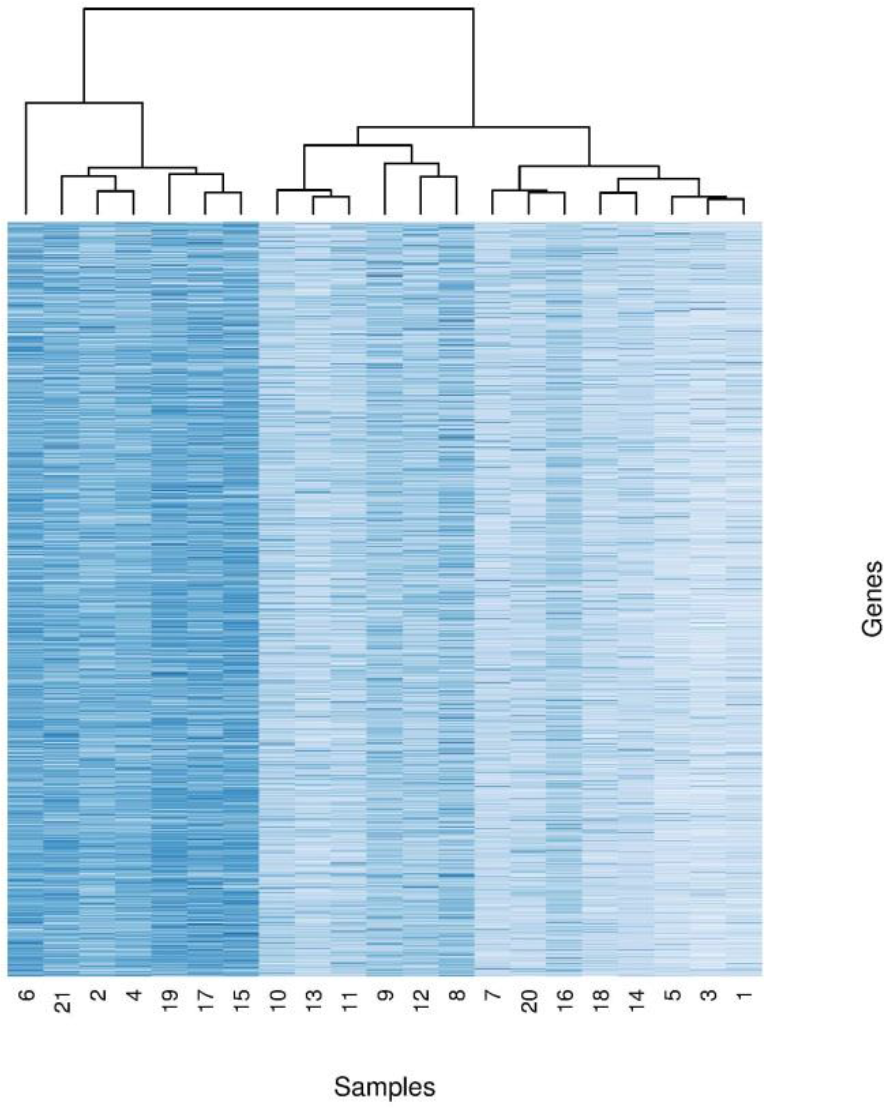
A heatmap showing the expression values for the genes in the Bottomly RNAseq data, showing a clear batch effect owing to the experiment number.

**Figure 5.**
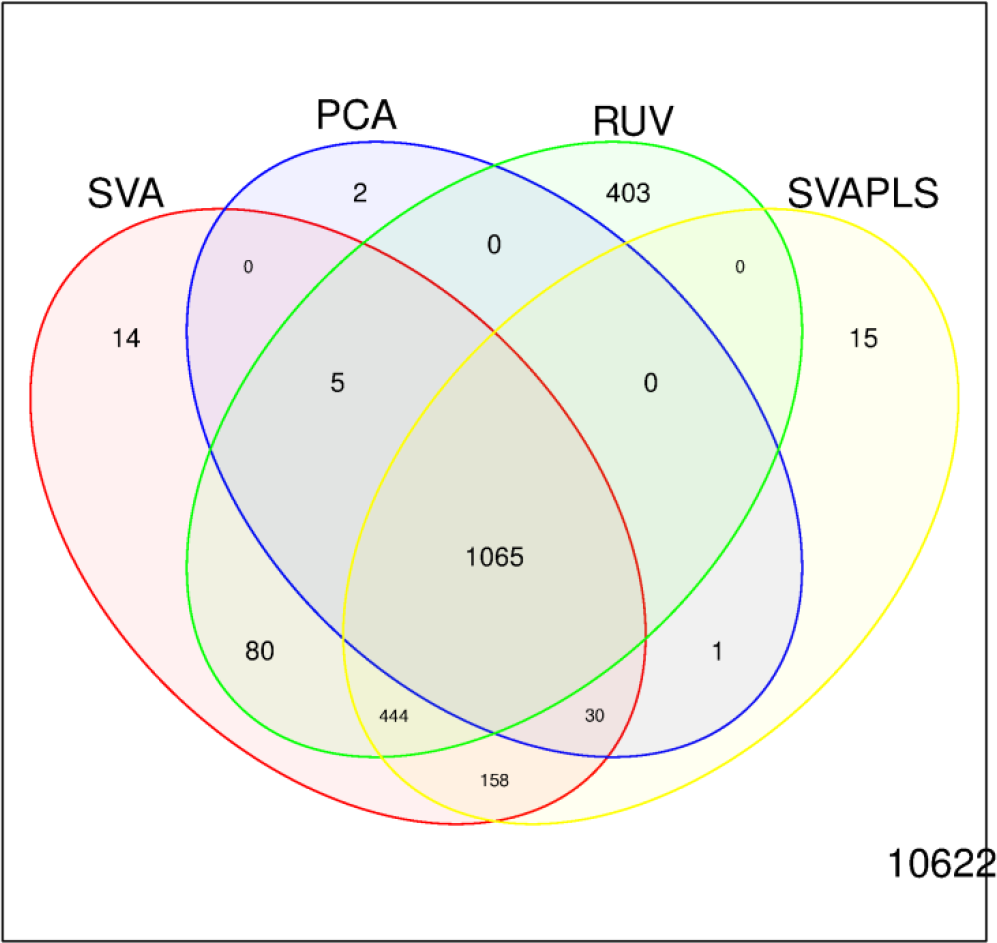
A Venn diagram of the differentially expressed genes detected by the six methods from an analysis of the Bottomly data.

**Figure 6.**
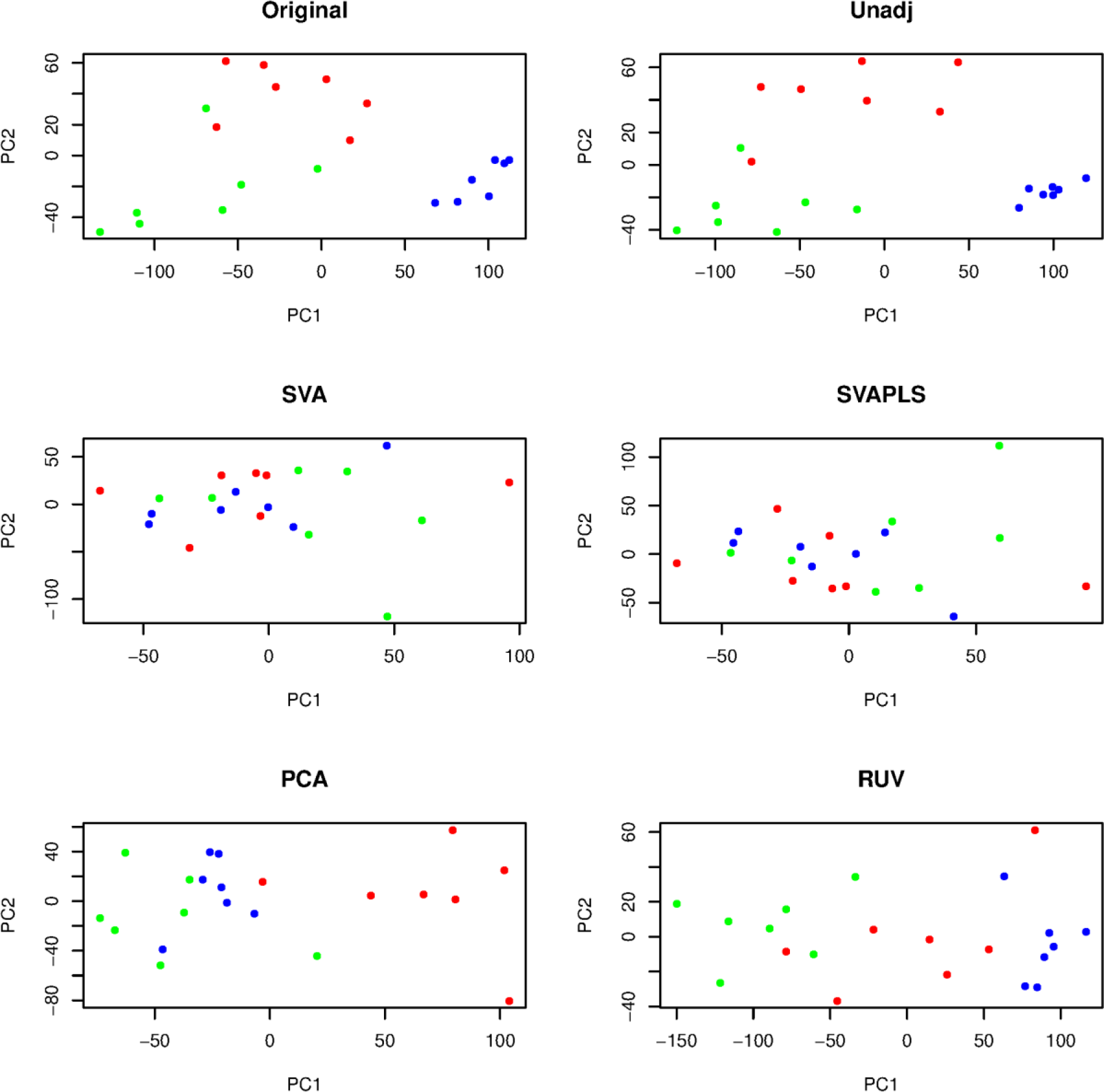
A scatterplot of the first and second principal components of the residual matrices obtained after removing the estimated signals of hidden variability in the data from the different methods. The red, green and blue dots represent the samples from experimental batches numbered 4, 6 and 7, respectively.

## Discussion

Latent variability in RNAseq gene expression data is attributable to several biological as well technical effects that are difficult to remove via standard normalization approaches. The complexity of the sequencing workflow along with the unknown biological profiles of the samples can potentially generate a wide array of hidden variables (confounders). Often, these variables impact the original pattern of differential gene expression between two groups of samples and introduce spurious signals of unwanted heterogeneity. As a result the commonly used tests based on simple linear models fail to detect a large number of true differentially expressed genes, while several genes are wrongly detected as false positives. I have introduced a novel method SVAPLSseq that identifies the signatures of different hidden effects in RNAseq gene expression data and corrects for them in order to enable a more powerful and accurate inference on the differentially expressed genes. The superior performance of the method has been validated by rigorous comparative studies with other available techniques, on both simulated as well as real-life RNAseq data. Both the studies demonstrate the impressive detection power and efficacy of hidden effect estimation by SVAPLSseq compared to the other methods. Moreover, the two variants of SVAPLSseq provide a wider scope of applying the method to different types of RNAseq data. In particular, the supervised version has been shown to be a much more powerful alternative than the competing methods when information on a set of control genes is available. Thus overall, the method provides a flexible and generalized framework to capture the effects of hidden variability in RNAseq gene expression data and adjust for them in order to potentially improve the detection power of differential expression analyses.

